# Multifaceted roles of rice ABA/stress-induced intrinsically disordered proteins in augmenting drought resistance

**DOI:** 10.1101/2023.06.22.546131

**Authors:** Meng-Chun Lin, I-Chieh Tseng, Ching-Lan Wang, Wen-Rong Hsiao, Yun-Jhih Shih, Wen-Dar Lin, Su-May Yu, Tuan-Hua David Ho

## Abstract

Water deficit stress causes devastating loss of crop yield worldwide. Improving crop drought resistance has become an urgent issue. Here we report that a group of abscisic acid (ABA)/drought stress-induced monocot-specific, intrinsically disordered, and highly proline-rich proteins, REPETITIVE PROLINE-RICH PROTEINS (RePRPs), play pivotal roles in drought resistance in rice seedlings. Rice ectopically expressing *RePRP*s outlived wild-type rice under extreme drought conditions primarily due to two underlying mechanisms. First, *RePRP* reduces water loss by decreasing stomata conductance in shoot. In addition, *RePRP* overexpression enhances the levels of extracellular water barriers such as lignin and suberin, primarily in the root vascular bundle. Several groups of genes involved in lignin biosynthesis, especially the wall-bound peroxidase responsible for the final assembly of the lignin network, were induced by *RePRP*. Second, overexpression of *RePRP* leads to lowered root osmotic potential. Root cell osmotic pressure was more negative in rice plants overexpressing *RePRP2* than wild-type plants, and the concentration of a key osmolyte, proline, was enhanced. Furthermore, the protein levels of two aquaporins that are important for drought stress tolerance were elevated. Hence, ABA/stress-induced *RePRP* expression leads to several beneficial traits of drought resistance, including lower water loss rate upon dehydration and higher root water use efficiency under drought conditions. This group of unique stress proteins may be an important target for technology development in enhancing drought stress resistance in cereals.

## Introduction

Plants experience various environmental stresses during their life cycles. Drought is one of the most detrimental stresses, causing severe loss of crop yield (Boyer 1982). As the climate has changed rapidly in recent decades, more regions have experienced inadequate water availability. Rice cultivars with high yield are usually cultivated in well-irrigated fields and are sensitive to water deficit. Thus, the development of drought-resistant crops is crucial to overcome the yield loss caused by water deficiency.

Researchers have been investigating physiological responses and molecular mechanisms for insights into improving innate crop resistance to drought. The current knowledge of abiotic stress is valuable for crop improvement with biotechnological approaches (Zhu 2002; Wang *et al*. 2003; Yamaguchi-Shinozaki and Shinozaki 2006; Seki *et al*. 2007). Water loss causes rapid changes in plant morphology and function, and plants use different strategies to overcome drought. One of the key features reflecting drought tolerance is the accumulation of osmolytes under water deficiency. The high concentration of osmolytes maintains cell turgor and elasticity with decreasing water potential of the environment (Morgan 1984). Sugars, sugar alcohols, betaines and proline are important osmolytes that accumulate during water stress to maintain growth and cell turgor (Voetberg and Sharp, 1991; Verslues and Sharp, 1999; Chen and Murata, 2002; Ashraf *et al*. 2007).

Mechanisms of drought avoidance maintain high water content in plant tissues in response to decreased environmental water. To escape drought conditions, plants have evolved a variety of physiological changes to minimize water loss and/or increase water uptake. Water-saving plants usually show reduced transpiration, whereas water-spending plants achieve high water content by increasing water uptake, mainly by increased rooting under drought (Basu *et al*. 2016). Numerous studies have shown that root growth is strongly inhibited under severe water stress (Westgate and Boyer, 1985; Sharp *et al*. 2004; Perkons *et al*. 2014.), but increased root length and density in the active root zone area improves water uptake and maintains high grain yield under water-deficit conditions (Sadok *et al*. 2011; Wasaya *et al*. 2018). The change in root morphology relies on altered cell wall integrity (Wu *et al*. 2000; Le Gall *et al*. 2015). Drought and high salinity lead to cell wall thickening and root tip swelling (Spollen *et al*. 2000; Ji *et al*. 2014; Li *et al*. 2014).

The drought-induced rapid root morphological changes are highly correlated with abscisic acid (ABA) content. Under drought conditions, ABA is produced to regulate a plethora of physiological responses (Shinozaki and Yamaguchi-Shinozaki, 2000; Finkelstein *et al*., 2002; Rabbani *et al*. 2003). The accumulation of ABA leads to signaling cascades that are required for proper induction of stress-response genes (Ingram and Bartels, 1996; Seki *et al*., 2002). Our previous research revealed that a group of rice ABA-inducible genes, *REPETITIVE PROLINE-RICH PROTEIN*s (*RePRP*s), highly expressed in the root elongation zone, were involved in regulating root growth and development. The rice genome has two small *RePRP* gene families; each encodes two almost identical proteins. Because of the very high content of proline (∼40%) throughout RePRPs, they are unlikely to form stable higher-order structures, hence being classified as intrinsically disordered proteins (Tseng *et al*., 2013). Increasing evidence has suggested that intrinsically disordered proteins with flexible structures such as *LATE EMBRYOGENESIS-ABUNDANT* (*LEA*) genes play pivotal roles in plant tolerance to water-deficit stress (Cuevas-Velazquez *et al*., 2021). RePRPs interact with cytoskeleton components (i.e., actins and tubulins) both *in vitro* and *in vivo*, which leads to the alteration of root tip development. Transgenic lines overexpressing *RePRP*s phenocopied wild-type (WT) rice (Tainung 67, TNG67) treated with ABA, showing short and swollen roots with more storage nutrients (Hsiao *et al*., 2020). In contrast, RNA-interference (RNAi) knockdown of *RePRP1* and *RePRP2* expression led to ABA insensitivity of root growth (Tseng *et al*. 2013). However, whether *RePRP*s are essential for rice drought tolerance remains to be addressed.

In this study, we investigated whether *RePRP*s are indeed crucial for rice survival under severe drought conditions. The beneficial traits of transgenic plants ectopically expressing *RePRP*s include several water conservation mechanisms in both roots and shoots, which facilitate the rice response to water deprivation.

## Results

### Rice *RePRP* overexpression enhances tolerance to dehydration stress

To assess whether *RePRPs* are required for drought tolerance, we generated a severe drought condition in our hydroponic system by treating rice seedlings with 30% polyethylene glycol (PEG). *RePRP*s are encoded by two small gene families, each containing two almost identical genes (Tseng *et al*., 2013). We used *RePRP2.1* (hereafter called *RePRP2*) for ectopic expression lines and *RePRP1* and *RePRP2* double RNAi lines throughout this research. After recovery from 30% PEG for 12 days, the mean survival of *RePRP2* overexpression lines (*RePRP2OX*) was 87.7±5.3%, with only 38.0±24.8% survival for WT plants (Fig. 1A and 1B, Fig. S1A). We also validated the increased drought survival of *RePRP2OX* lines in a large number of additional independent experiments with slightly less stressful 25% PEG (Fig. S1B). In contrast, mean drought survival rates for RNAi lines that knocked down both *RePRP1* and *RePRP2* (*RePRP*-RNAi) were 14.0±20.7% to 24.5±29.2%, which is in general more sensitive to drought as compared with the WT (Fig. 1B). Furthermore, the *RePRP2OX* lines showed better performance under dried soil conditions as compared with both WT and *RePRP*-RNAi lines (Fig. 1C, Fig. S1C). Because constitutive overexpression may not reflect the actual stress responses mediated by *RePRP*s, we generated transgenic *RePRP2OX* lines driven by a synthetic ABA-inducible promoter (*3X ABRC321*), which is mainly expressed in roots under abiotic stress conditions (Chen *et al*. 2015). Induced overexpression of *RePRP2* (*ABRC:RePRP2*) also significantly increased rice survival with 30% PEG treatment as compared with the WT (Fig. 1D and 1E). Moreover, the effects of stress on root length and morphology seen in the constitutive *RePRP2OX* lines were not observed in the *ABRC:RePRP2* plants, so the drought-induced expression of *RePRP2* retained the benefits of drought tolerance along with normal root growth in the absence of abiotic stress (Fig. 1A and 1D). These data suggest that *RePRP2* is essential for maintaining rice growth and development under drought.

**Figure 1.**
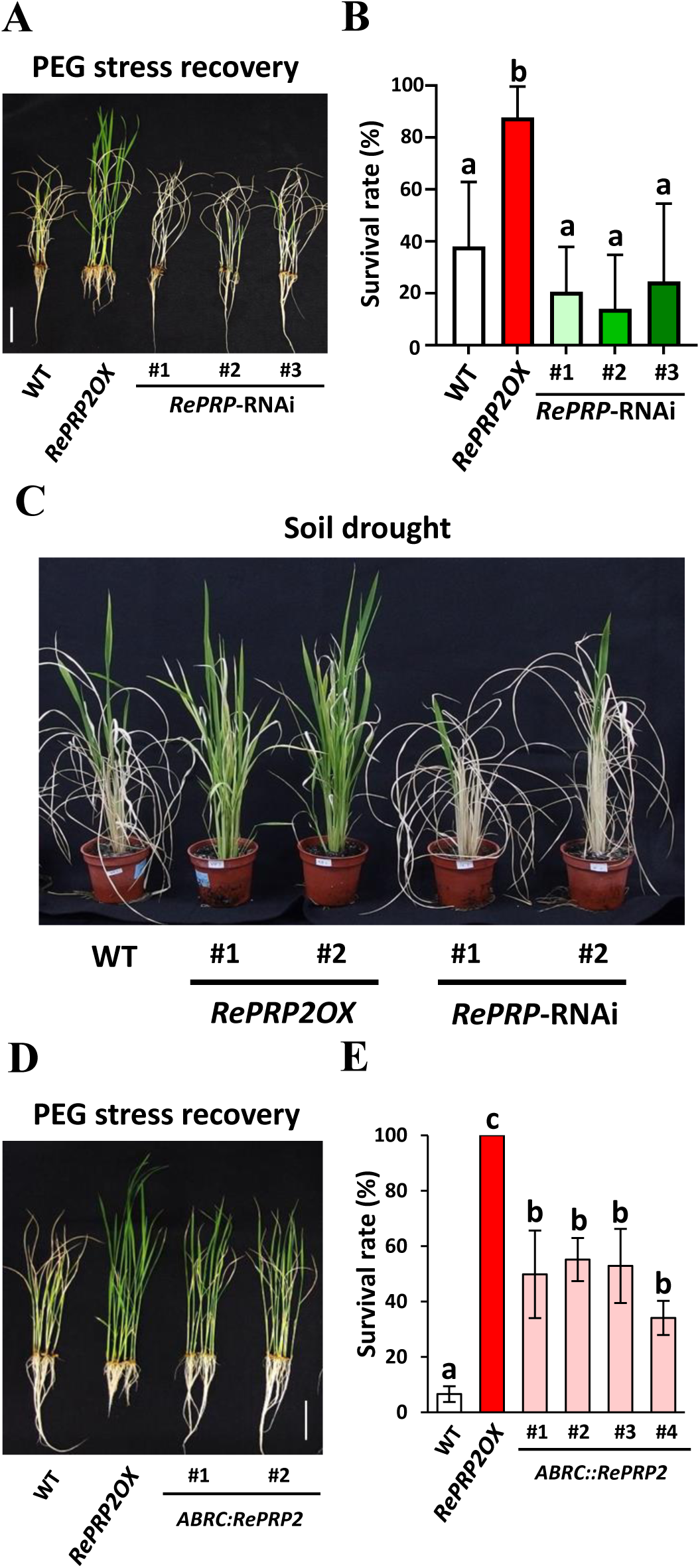
*RePRP2* is essential for drought survival. **A B** *RePRP2* overexpression lines (*RePRP2OX*) showed increased survival rate and RNAi lines reduced survival after recovering from extreme drought stress (30% PEG). Scale = 5 cm. Three-week-old rice plants grown in ½ Kimura solution were treated with 30% PEG for 20 h, followed by recovery in ½ Kimura solution. The survival rate was calculated on day 15 after recovery. Data were analyzed by one-way ANOVA with Tukey’s multiple comparison test. Data are shown as means ± SD from four biological repeats, with 30 seedlings per line in each repeat. Different superscript letters (a, b) indicate significant mean difference (p < 0.05). **C** Soil culture results of resistance to drought stress for *RePRP2OX* lines and *RePRP*-RNAi lines. One-month-old soil-grown rice plants were withheld from watering for 4 days before re-watering. Image was taken at day 15 after recovery. **D E** Stress-inducible promoter-driven *RePRP2OX* lines (*ABRC::RePRP2*) showed increased drought survival compared to the wild type (WT). Scale = 5 cm. Data are mean ± SD from 30 individuals. At least three independent experiments were conducted and all showed similar results. PEG treatment procedures and analysis are as in Fig. 1A/B. Data were analyzed by one-way ANOVA with Tukey’s multiple comparison test. Different superscript letters indicate significant mean difference (p < 0.05).

### *RePRP2* reduces rice water loss and increases root water use efficiency

Given that *RePRP2* overexpression lines are highly resistant to drought stress and its transcript level is highly induced by ABA treatment (Fig. S2A), we further investigated the detailed mechanisms of *RePRP*-mediated drought resistance. Our dehydration assays showed that *RePRP2OX* plants still maintained 45% to 55% of water after 2 h of dehydration, with only 30% of water left in WT and *RePRP*-RNAi plants. (Fig. 2A–2B). Transpiration is the main driving force of plant water transport and is highly related to stomata development and conductance. Rice plants show decreased stomata density and conductance in response to water deficiency, accompanied by reduced transpiration, which prevents plants from excessive water loss (Caine *et al.,* 2019). A previous report suggested that although *RePRP2* was mainly expressed in root, it was also expressed in shoot (Tseng *et al*., 2013). To assess the rate of root water loss, we have conducted the dehydration assay with leaves covered by Vaseline to block water loss from transpiration. Our results showed that WT still contains 50% of water after 8h of dehydration, while *RePRP2OX* lines still showed reduced root water loss; the rate of water loss in *RePRP*-RNAi was comparable to WT (Fig. 2C – 2D). Our qRT-PCR analyses also indicated that *RePRP2* can be induced by ABA in the shoot, but the transcript level was lower than in the root (Fig. S2B and S2C). Rice transformed with an *RePRP2* promoter-driven *GUS* gene cassette (*ProRePRP2.1::GUS*) exhibited β-glucuronidase (GUS) staining signals in guard cells (Fig. 2E). On scanning electron microscopy, *RePRP2OX* lines showed slightly higher stomata density than WT and *RePRP*-RNAi lines (Fig. 2F–2I). The size of *RePRP2OX* stomata was significantly reduced as compared with WT and *RePRP*-RNAi leaves (Fig. 2I), in agreement with the lower stomata conductance in *RePRP2OX* lines (Fig. 2K). In summary, *RePRP2* is sufficient for reducing stomatal conductance and water loss under drought.

**Figure 2.**
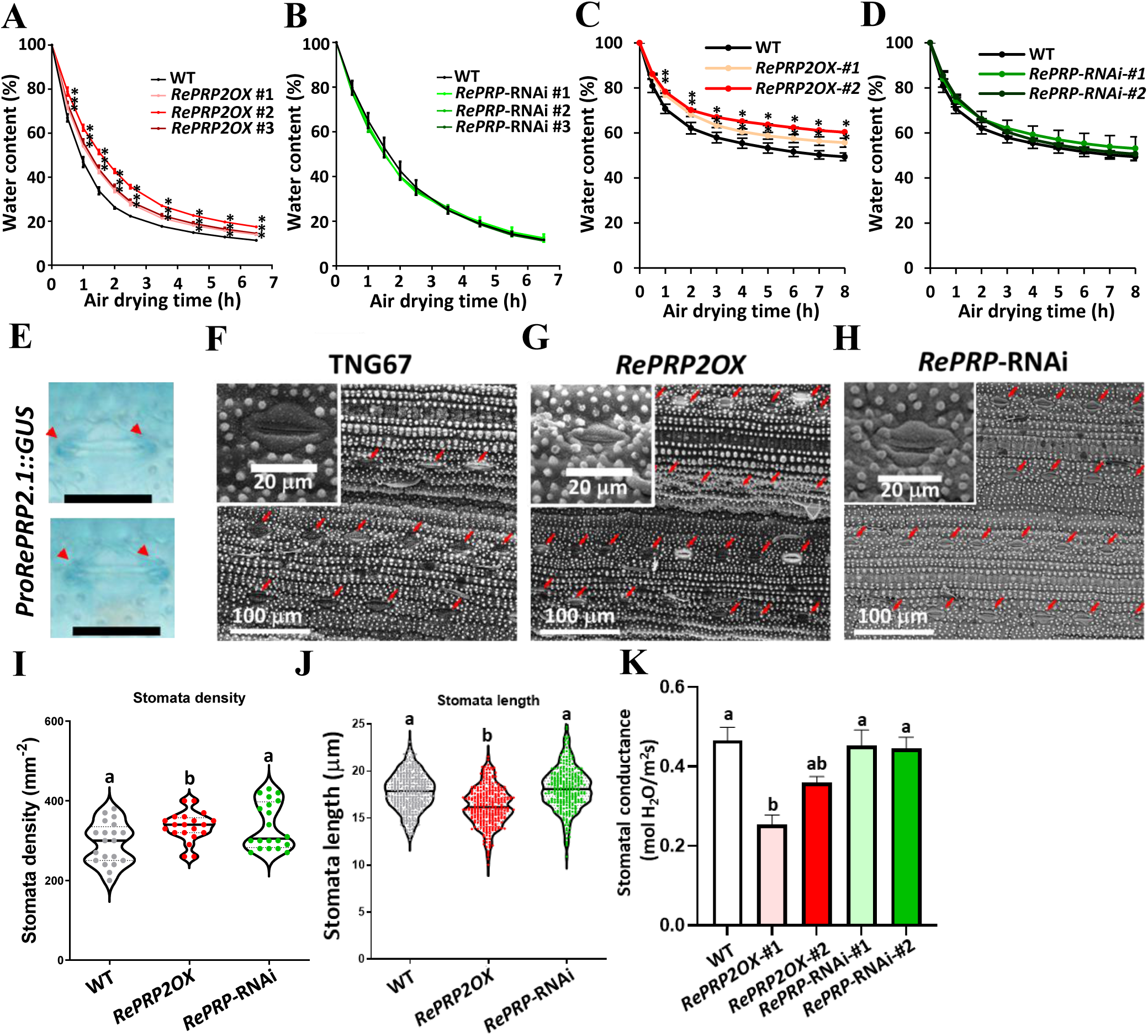
*RePRP2* reduces water loss during seedling dehydration. **A** *RePRP2OX* lines retain more water under dehydration. Ten-day-old WT and *RePRP2OX* plants dehydrated in air. The fresh weight was measured at different times to calculate relative water content (%) remaining in plants. Asterisks indicate significant difference to WT at the same time point by Student *t* test (p < 0.05). **B** *RePRP*-RNAi lines showed comparable water loss rate as the WT. Data are mean ± SD from 15 individual plants. At least three independent experiments were conducted and all showed similar results. **C D** *RePRP2* reduced root water loss. The shoots of 10-d-old seedlings were covered with vaseline to block water loss from transpiration; the fresh weight before and after vaseline spreading was recorded. The plants were then dehydrated in air. Data are mean ± SD from 10 seedlings. The fresh weight was measured at different times to calculate relative water content (%) remaining in plants. Asterisks indicate significant difference from the WT at the same time point by Student *t* test (p < 0.05). **E** *RePRP2* was expressed at guard cells. β-glucuronidase activity assay was used to visualize the localization of *RePRP2.1* expression. Ten-day-old seedlings of *ProRePRP2.1::GUS* grown in ½ Kimura solution were used for GUS activity staining. Scale: 25 μm. Red arrows indicate sites of GUS signal accumulation. Two typical stomata are shown. **F G H** SEM imaging showing *RePRP2OX* leaves with increased stomata density. The upper epidermis from the middle part of the leaves of 14-day-old seedling were used for observations. Red arrowheads indicate the positions of stomata. **I** *RePRP2OX* lines show increased stomata density, whereas *RePRP*-RNAi lines show comparable stomata density to the WT. Violin plots of the overall distribution of stomata density calculated from 10 images per line. At least two biological replicates were conducted and all showed similar results. Solid horizontal lines indicate mean stomata length, whereas dashed horizontal lines indicate interquartile range. Data were analyzed by one-way ANOVA with Tukey’s multiple comparison test. Different superscript letters indicate significant mean difference (p < 0.05). **J** *RePRP2OX* lines show reduced stomata size. Violin plots of the overall distribution of stomata length calculated from 10 images per line. Horizontal lines indicate mean stomata length. At least two biological replicates were conducted and all showed similar results. Data were analyzed by one-way ANOVA with Tukey’s multiple comparison test. Different superscript letters indicate significant mean difference (p < 0.05). **K** *RePRP2OX* lines show decreased stomatal conductance, whereas *RePRP*-RNAi lines show comparable stomatal conductance to WT plants. Two-month-old rice plants grown in soil were used for measuring stomatal conductance. Data are mean ± SD from 12 individual plants. Data were analyzed by one-way ANOVA with Tukey’s multiple comparison test. Different superscript letters indicate significant mean differences (p < 0.05).

In addition to reduced water loss under drought, *RePRP2OX* lines had better recovery after drought (Fig. 1A–1E). Considering that *RePRP2* is highly expressed in root (Fig. S2B), we reasoned that the induction of *RePRP2* may facilitate root water uptake under water deficit. To study the efficiency of root water uptake, we treated rice seedlings with mild drought, then covered the leaves with Vaseline to block transpiration. The Vaseline-covered plants were incubated with distilled water to evaluate the fresh weight recovery by root water uptake. *RePRP2OX* lines regained up to 75% of their original fresh weight after 6h of rehydration, which is significantly higher than that of WT and *RePRP*-RNAi lines, suggesting that *RePRP2* facilitates root water uptake after re-hydration (Fig. 3A). *RePRP2OX* plants also consumed less water than the WT under normal conditions, to a level similar to PEG-treated WT plants (Fig. 3B and 3C). These results implied that *RePRP2* is likely involved in regulating water use efficiency (WUE). WUE is a critical indicator of crop production, especially under water deficit (Condon *et al*. 2004; Karaba *et al*. 2007). Our results showed that the root WUE was higher for *RePRP2OX* than WT plants and was comparable to WT plants treated with 10% PEG (Fig. 3D and Fig. S3A). However, shoot WUE did not differ between WT and *RePRP2OX* plants (Fig. 3E and Fig. S3B). The water use and WUE in *RePRP*-RNAi lines was comparable to that of the WT (Fig. S3C and S3D). In summary, *RePRP2OX* lines used less water to increase root biomass, resembling WT seedlings treated with mild osmotic stress.

**Figure 3.**
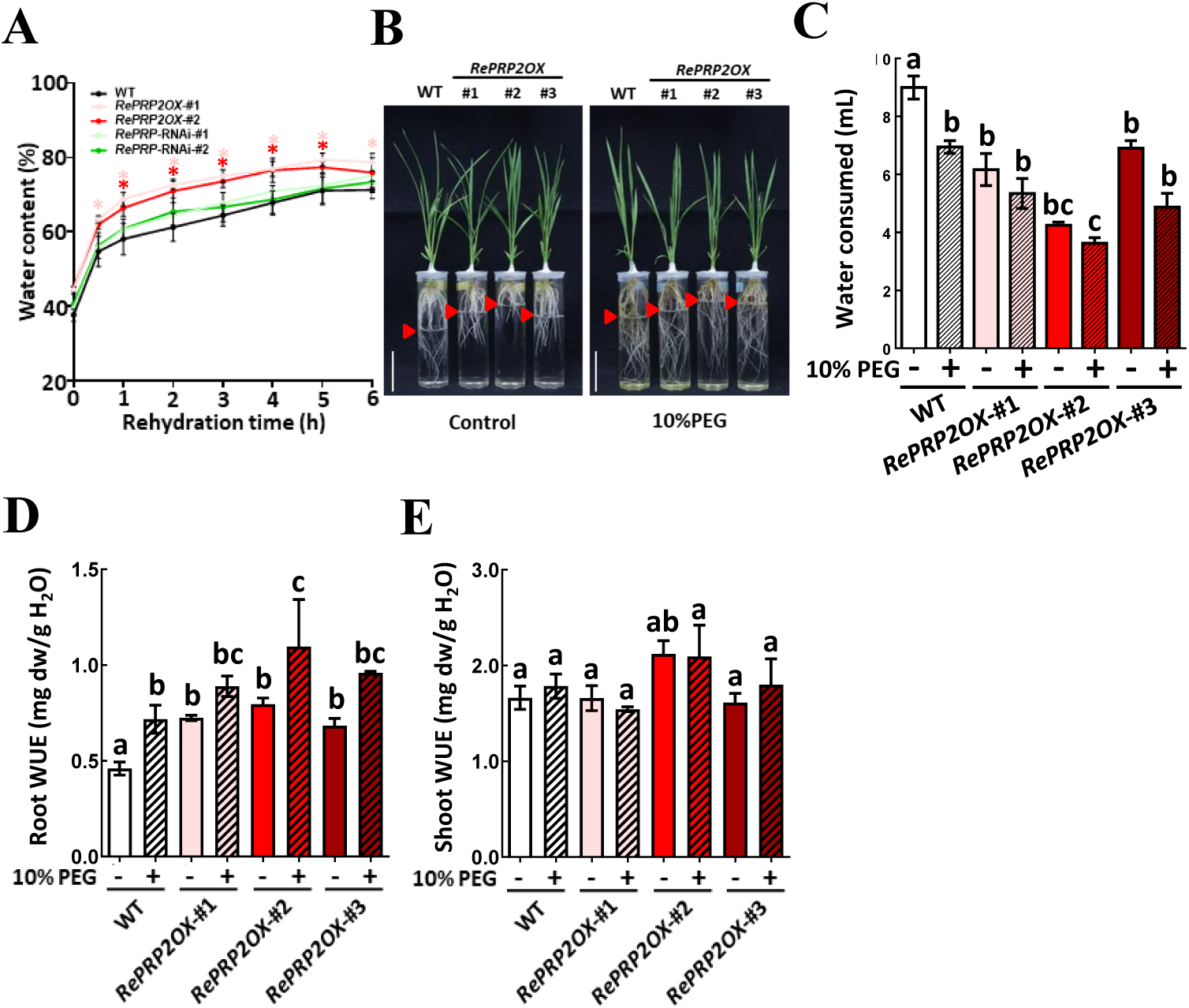
*RePRPs* enhance root water uptake and water use efficiency. **A** *RePRP2OX* lines showed increased water uptake after drought. Ten-day-old rice seedlings were dried in air for 3 h and leaves were sealed with vaseline to block transpiration; the fresh weight before and after vaseline spreading was recorded. The plants were put into 100 mL distilled water and fresh weight was measured at each time point. Data are mean ± SD from 10 seedlings. 100% water content was defined as fresh weight prior to dehydration. The tested plants were then dried in 50°C oven for 2 days and the plant dry weight were recorded and defined as 0% of water content. Colored asterisks indicate significant difference from the WT at the same time point by Student *t* test (p < 0.05). **B C** *RePRP2OX* lines use less water. The amount of water consumed in air-tight test tubes of 10-day-old WT and *RePRP2OX* lines was measured after 1 week of incubation. Each tube contains four seedlings of the WT or *RePRP2OX* lines. Arrows indicate the position of the water surface of each tube. Data were analyzed by one-way ANOVA with Tukey’s multiple comparison test. Different superscript letters indicate significant mean difference (p < 0.05). **D** Root water use efficiency (WUE) was increased in *RePRP2OX* lines. Data are mean ± SD from 12 individual plants. Data were analyzed by one-way ANOVA with Tukey’s multiple comparison test. Different superscript letters indicate significant mean difference (p < 0.05). **E** Shoot WUE was not significantly changed in *RePRP2OX* lines. Data were analyzed by one-way ANOVA with Tukey’s multiple comparison test. Different superscript letters indicate significant mean difference (p < 0.05).

### *RePRP2* is required for ABA-mediated induction of some drought responsive genes

Our previous studies showed that *RePRP2* is mainly expressed in the elongation zone of root and *RePRP2OX* showed swollen root phenotype similar to WT roots treated with ABA (Tseng *et al*., 2013). Thus, we collected root tips (1 cm from the top of roots) from WT, *RePRP2OX* and *RePRP*-RNAi plants and extracted total RNA for transcriptomic analyses. The genes with greater than 2-fold change in expression as compared with the TNG67 WT under control condition were defined as differentially regulated genes. Our transcriptomic analyses showed that about 11.6% of the expressed genes were differentially upregulated by ABA treatment or *RePRP2* overexpression (1728 of 20781 genes), with 152 genes commonly upregulated by both ABA and *RePRP2OX* (Table S1). As shown in the leftmost blocks of Figure S3, the red color indicates that many key transcription factors (TFs) involved in stress responses were highly upregulated by ABA. The transcript levels of these TFs were also elevated in *RePRP2OX* lines in the absence of ABA. In contrast, the ABA induction of these TFs were less effective in *RePRP*-RNAi lines as compared with ABA-treated WT (Fig. S4, Table S2). Our transcriptomic data suggests that *RePRP* is sufficient and necessary for the ABA induction of these TFs. Most of the ABA- and *RePRP2*-dependent TF families belong to *WRKY*, *NAM, ATAF and CUC* (*NAC*), *ETHYLENE-RESPONSIVE FACTOR* (*ERF*), *HEAT SHOCK FACTOR* (*HSF*) and *HOMEOBOX* (*HOX*) families (Fig. 4A and Fig. S4 and Fig. S5). TFs that belong to these families include *DROUGHT RESPONSIVE ZINC FINGER PROTEIN 1* (*DRZ1*), *WRKY21*, *WRKY76*, *NAC066*, *NAC095, HSF15* and group VII *ERF*s, reported to enhance abiotic stress tolerance. (Yuan *et al*., 2018; Licausi *et al*., 2013; Hoang *et al*., 2019; Shen *et al*., 2009; Sun *et al*., 2015; Wang *et al*., 2020; Yokotani *et al*., 2013). We also validated the expression patterns of *HSF15* and *NAC066* by qRT-PCR. Consistent with our transcriptomic analyses, ABA and *RePRP2OX* increased the level of all the examined TFs, whereas ABA induction of *HSF15* and *NAC066* was reduced in the *RePRP*-RNAi line (Fig. 4B–4C; Fig. S5).

**Figure 4.**
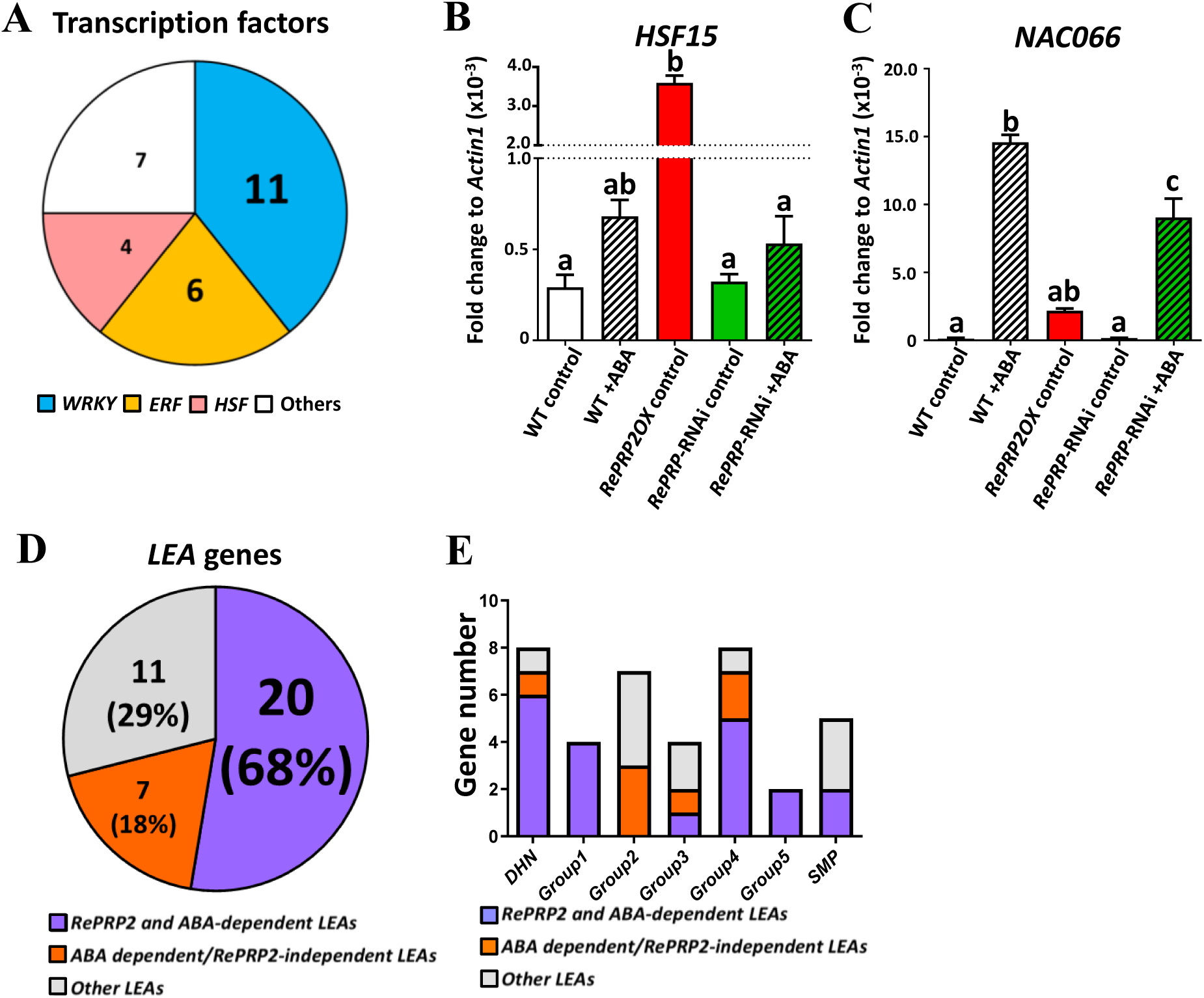
*RePRP2* is necessary and sufficient for ABA-mediated induction of drought-responsive genes. **A** Stress-responsive transcription factors (TFs) that were highly activated by ABA application and *RePRP2* overexpression. Numbers in the pie chart indicate genes within the gene family showing differential induction in response to ABA application and *RePRP2* overexpression. TFs with more than 2-fold change by ABA application and *RePRP2* overexpression compared to TNG67 control were defined as differentially induced TFs. **B C** qRT-PCR validation of ABA and *RePRP2*-dependent TFs identified in the root tip transcriptome. *HSF15* and *NAC066* were chosen as target genes for validation. Ten-day-old seedlings were treated with or without ABA (2 μM) for 96 h before root tissue collection (1 cm from root tip) and total RNA extraction. Data are mean ± SD from three technical repeats, with *Actin1* as an internal control. Data were analyzed by one-way ANOVA with Tukey’s multiple comparison test. Different superscript letters indicate significant mean difference (p < 0.05). **D** ABA-induced *LEA* genes were highly upregulated in *RePRP2OX* lines under control conditions. Numbers indicate *LEA* genes responsive to ABA and/or *RePRP2* overexpression. All expressed *LEA*s in the transcriptome were used to calculate the gene numbers in the pie chart. *LEA*s with more than 2-fold change by ABA application and *RePRP2* overexpression compared to TNG67 control were defined as *RePRP2* and ABA-dependent *LEA*s. *LEA*s with more than 2-fold change after ABA application but less than 2-fold change with *RePRP2* overexpression were defined as ABA dependent/*RePRP2*-independent *LEA*s. *LEA*s with less than 2-fold change with both ABA application and *RePRP2* overexpression were defined as other *LEA*s. **E** Statistics of expressed *LEA*s in rice root tip transcriptomes. *RePRP2* and ABA-dependent *LEA*s were enriched in the *DEHYDRIN* (*DHN*), group 1, group 4 and group 5 *LEAs*. *SMP*: seed maturation protein.

We also observed that the expression of abiotic-stress-responsive *RESPONSIVE TO ABA GENE 16A* (*Rab16A*) was highly dependent on the presence of *RePRP2* (Fig. S5). *Rab16A* belongs to the *LEA* gene family, suggested to be essential for rice drought and stress tolerance (Wang *et al*., 2007, Xiao *et al*., 2007). Our transcriptomic analyses revealed that *RePRP2* was necessary and sufficient for the ABA-mediated upregulation of at least 20 *LEA* genes (Fig. 4D, Fig. S6, Table S3). *LEA* genes were classified into six groups based on their deduced amino acid sequences (Wang et al., 2007). The ABA- and *RePRP*-dependent *LEA*s were specifically enriched in *DEHYDRIN* (*DHN*), group_1, group_4 and group_5 *LEAs* (Fig. 4E). The *RePRP*-dependent ABA induction of stress-responsive TFs and *LEA* genes suggested that *RePRP*s are crucial regulators of ABA-mediated drought responses.

### *RePRP2* is essential for stress-enhanced root cell-wall lignification and suberization formation

The short and swollen root morphology and cell wall cellulose microfibril orientation of *RePRP2OX* lines resembled ABA-treated WT roots (Tseng *et al*., 2013; Hsiao *et al*., 2020). Several *NAC* TFs, such as *NAC066* and *NAC095*, potentially involved in cell wall development (Shen *et al*., 2009; Sun *et al*., 2015), were also upregulated by *RePRP2OX* (Fig. S4). We reasoned that the ABA-mediated *RePRP* induction may be related to changes in cell wall structure/function in response to stress. Our Gene Ontology (GO) and Kyoto Encyclopedia of Genes and Genomes (KEGG) pathway analyses revealed that functions of genes commonly upregulated by ABA and *RePRP2* overexpression were enriched in the ABA-activated signaling pathway (p = 2.9 x 10^-3^, Fisher’s exact test) and cell wall macromolecule catabolism process (p=1.1 x 10^-3^, Fisher’s exact test) (Table S4). In support of these discoveries, the genes involved in cell wall metabolism and phenylalanine metabolism, especially the cell-wall-bound peroxidases (*POX*s) responsible for lignin biosynthesis, were enriched among ABA and *RePRP2OX* co-regulated genes (Table S4). *POX*s are the only genes that showed both ABA and *RePRP2* dependence in the lignin biosynthesis pathway (Fig. 5A and Table S5 and S6). Our qRT-PCR results also validated that the expression of *POX51-like*, *OsPRX86* and *OsPRX90* highly depended on ABA application and *RePRP2* overexpression (Fig. 5B–5D). Cell wall-bound POXs are responsible for catalyzing lignin and suberin polymerization by transferring electrons from H_2_O_2_ to lignin and suberin monomers (Douglas, 1996; Moura *et al*. 2010; Wolf and Höfte, 2014; Graça, 2015; Liu *et al*., 2018). Consistent with our transcriptomic analyses, *RePRP2OX* lines showed increased root H_2_O_2_ accumuolation and cell-wall–bound POX activity (Fig. 5E and 5F). Our suberin fluorescence staining showed that *RePRP2OX* lines emitted higher suberin fluorescence than the WT and *RePRP*-RNAi lines under control conditions (Fig. S7). The application of ABA led to increased fluorescence signals at xylem, but this increase was attenuated in *RePRP*-RNAi lines (Fig. S8). The results suggest that *RePRP2* is responsible for ABA-mediated suberin accumulation, especially in the root vascular tissue. A similar phenomenon of lignin accumulation was observed in *RePRP2OX* lines, as revealed by fluorescence staining (Fig. S9). ABA application caused specific accumulation of lignin in only the xylem of the WT and *RePRP*-RNAi lines but could not further increase the fluorescent signals in the *RePRP2OX* line, which suggests that the effect of *RePRP2OX* was overshadowed by the effects of ABA (Fig. S9, S10). We also extracted root cell walls with acetyl bromide and validated a ∼20% increase of root tip lignin in *RePRP2OX* versus WT and *RePRP*-RNAi lines, resembling WT root tips treated with ABA (Fig. 5G). Our discoveries suggest that *RePRP* is sufficient for the accumulation of root suberin and lignin, possibly by enhancing cell wall-bound POX activity (Fig. 5 and Fig S7–S10). The increase in root suberin is critical for regulating water loss, and lignin deposition is essential for controlling root permeability (Liu *et al*., 2018; de Silva *et al*., 2021; Reyt *et al*., 2021). The increased root lignin/suberin content in roots, together with reduced transpiration in shoots, may have reduced water loss of *RePRP2OX* under drought.

**Figure 5.**
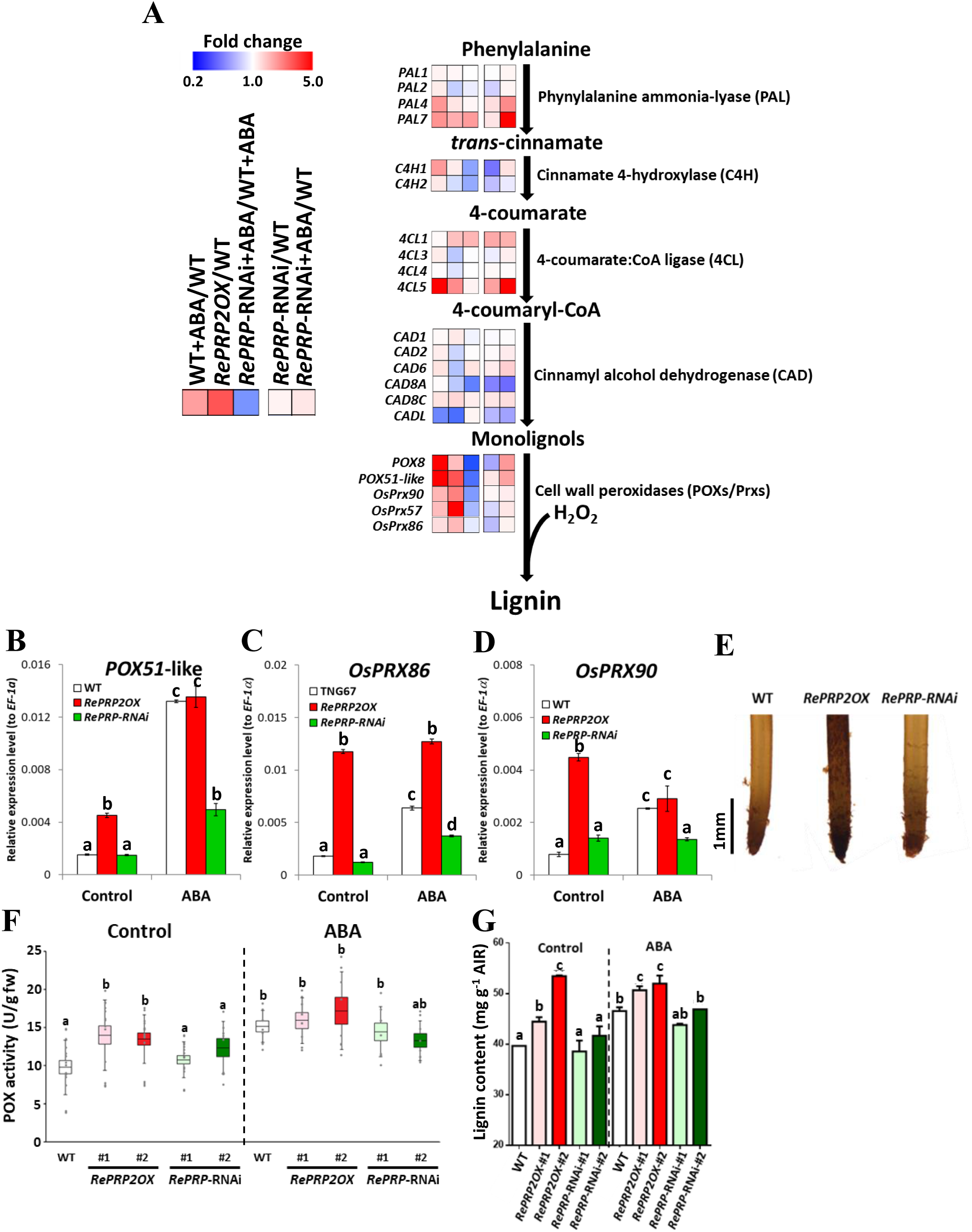
*RePRP*s are required for suberin and lignin formation in root. **A** Transcriptomic analyses showing upregulation of the critical enzymes involved in the lignin biosynthesis pathway by *RePRP2* overexpression and ABA (2 μM) treatment. Ten-day-old seedlings were treated with or without ABA (2 μM) for 96 h before root tip collection and total RNA extraction. The relative mRNA levels of lignin biosynthesis genes expressed in the root transcriptome are shown. *PAL*: phenylalanine aminolyase; *C4H*, cinnamate 4-hydroxylase; *4CL*, 4-coumarate:CoA ligase; *CAD*, cinnamyl alcohol dehydrogenase; *POX/Prx*, cell wall peroxidases reported to be involved in lignin biosynthesis. Data are shown as heat maps based on the mean fold change from three biological repeats. The leftmost block of each gene indicates the fold change in expression of WT plants treated with ABA to the WT control, and the second block indicates the fold change in expression of *RePRP2OX* control plants to the WT control. The third block indicates the fold change in expression of *RePRP*-RNAi lines treated with ABA to the WT treated with ABA. The “red-red-blue” patterns indicate that *RePRP*s are necessary and sufficient for ABA induction of these genes. The fourth and fifth blocks show fold change in expression of *RePRP*-RNAi lines to the WT control, with the absence or presence of ABA, respectively. **B C D** qRT-PCR validated that *RePRP2* is essential for cell wall-bound peroxidase expression. *RePRP2OX* showed elevated levels of *POX51*-like, *OsPRX86* and *OsPRX90*. Data are mean ± SD from three technical repeats, with *EF-1α* as an internal control. Data were analyzed by one-way ANOVA with Tukey’s multiple comparison test. Different superscript letters indicate significant mean difference (p < 0.05). **E** *RePRP2OX* lines show increased H_2_O_2_ content in roots. DAB staining was used to visualize the accumulation of H_2_O_2_. Scale: 1 mm. **F** *RePRP2OX* lines show elevated root cell wall-bound peroxidase activity. Ten-day-old rice seedlings were treated with or without 2 μM ABA for an additional 24 h. Data are mean ± SD. One unit of POX activity was defined as 1 mmol tetraguaiacol produced per minute. At least three biological replicates were conducted and all showed similar results. Horizontal lines, boxes and whiskers indicate mean, SE and SD, respectively. Data were analyzed by one-way ANOVA with Tukey’s multiple comparison test. Different superscript letters indicate significant mean difference (p < 0.05). **G** *RePRP2* is required for root tip lignin accumulation. Lignin content in root tips (1 cm) measured in WT and transgenic plants. Ten-day-old rice seedlings were treated with or without 2 μM ABA for an additional 4 days. Data are mean ± SD from three technical repeats. At least three biological replicates were conducted and all showed similar results. Data were analyzed by one-way ANOVA with Tukey’s multiple comparison test. Different superscript letters indicate significant mean difference (p < 0.05).

### *RePRP2* lowers root osmotic potential and enhances water transportation

*RePRP2OX* lines showed increased root water uptake under drought (Fig. 3A). Plants reduce osmotic potential by increasing soluble sugars, amino acids and ionic content in the cytosol to promote water uptake from dehydrated soil. Metal ions and osmolytes such as sugars, sugar alcohols, betaines and proline are important chemicals that accumulate during water stress (Voetberg and Sharp, 1991; Verslues and Sharp, 1999; Chen and Murata, 2002; Ashraf *et al*., 2007). Our *RePRP2OX* lines showed more negative root osmotic potential (-0.51 to -0.59 MPa) than did WT seedlings (-0.49 MPa), reaching a level similar to WT plants treated with 15% PEG (-0.54 MPa), thus allowing root water uptake even under osmotic stress (Fig. 6A). Our inductively coupled plasma optical emission spectrometry (ICP-OES) showed that *RePRP2OX* roots had comparable concentrations of potassium, sodium, phosphorus, iron and calcium to WT and *RePRP*-RNAi lines, so metal ions do not appear to be the main source of low osmotic potential in *RePRP2OX* roots (Fig. S11). We then focused on other osmolytes that may also cause decreased osmotic potential. Proline content was significantly increased in *RePRP2OX* roots to a level similar to WT plants treated with ABA. *RePRP*-RNAi lines showed comparable proline content to WT only under both control and ABA-treated conditions (Fig. 6B). Furthermore, the transcript levels of pyrroline-5-carboxylate synthase 1 (*P5CS1*) and pyrroline-5-carboxylate reductase (*P5CR*) were elevated in *RePRP2OX* roots under control conditions, similar to WT roots treated with ABA. However, the levels of *P5CS* and *P5CR* under ABA were slightly repressed in *RePRP*-RNAi lines (Fig. S12A and S12B; Table S8). The gene responsible for proline degradation, proline dehydrogenase (*ProDH*), was also increased in level in *RePRP2OX* under control conditions, whereas ABA treatment severely decreased the level of *ProDH* in all lines tested (Fig. S12C). Our results suggest that the lowering of root osmotic potential in *RePRP2OX* lines is partly due to increased root proline content.

**Figure 6.**
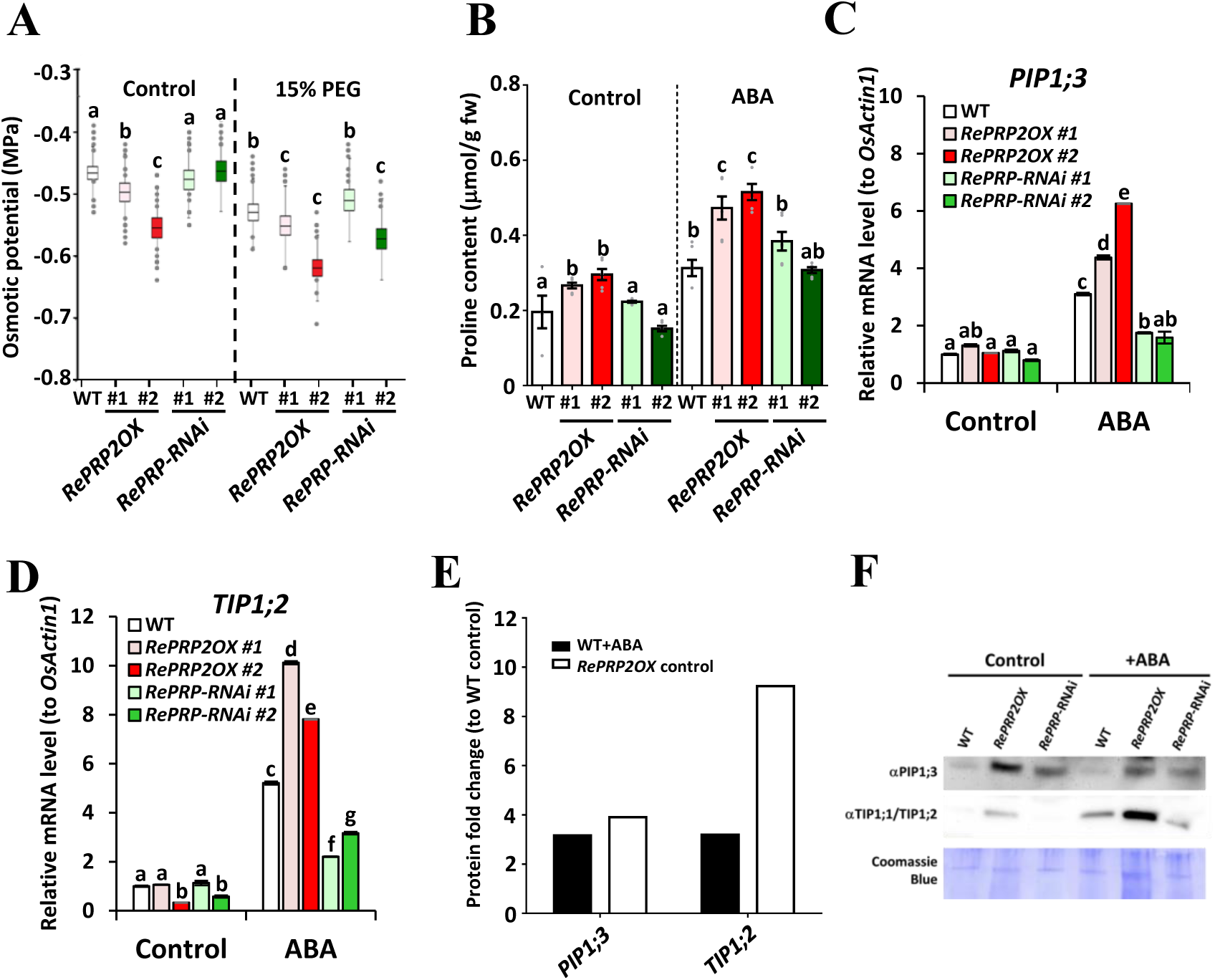
*RePRP*s lower root osmotic potential and increase aquaporin expression. **A** *RePRP2OX* lines showed more negative osmotic potential as compared with the WT and *RePRP*-RNAi lines under control conditions. Seedlings grown in 15% PEG treatment showed a further decrease of osmotic potential. The horizontal line, boxes and whiskers indicate mean ± SE and mean ± SD, respectively. Data were analyzed by one-way ANOVA with Tukey’s multiple comparison test. Different superscript letters indicate significant mean difference (p < 0.05). **B** ABA application and *RePRP2* overexpression increased root proline content. Ten-day-old rice seedlings were treated with or without 2 μM ABA for an additional 1 day. Data are mean ± SD. At least three biologal replicates were conducted and all showed similar results. Data were analyzed by one-way ANOVA with Tukey’s multiple comparison test. Different superscript letters indicate significant mean difference (p < 0.05). **C D** The transcript levels of aquaporins were elevated by ABA and *RePRP2* overexpression. Time-course qRT-PCR results of plasma membrane- (*PIP1;3*) and tonoplast-type aquaporin (*TIP1;2*) after 96 h of ABA treatment in WT, *RePRP2OX* lines, and *RePRP2OX* lines. Ten-day-old rice seedlings were treated with or without 2 μM of ABA for an additional 96 h. Whole root tissues were collected for total RNA extraction and qRT-PCR. Relative expression compared to the internal control *OsActin1* is shown as mean ± SD. At least three biological replicates were conducted and all showed similar results. Data were analyzed by one-way ANOVA with Tukey’s multiple comparison test. Different superscript letters indicate significant mean difference (p < 0.05). **E** Proteomic analysis of ABA application and *RePRP2* overexpression elevating levels of PIP1;3 and TIP1;2. The protein fold change was based on the abundance ratio between samples from Mascot search. **F** Western blot analysis of PIP1;3 and TIP1;1/TIP1;2 levels under control conditions in WT, *RePRP2OX* and *RePRP*-RNAi lines after ABA application. Ten-day-old seedlings grown in ½ Kimura solution were treated with or without 2 μM of ABA for an additional 96 h. An amount of 5 μg of root protein extracts was added to each well and Coomassie Blue G250-stained membrane was the loading control.

Aquaporins (AQPs) are important water transporters in plant tissues and play pivotal roles in water transportation in response to drought (Afzal *et al*., 2016). AQP-mediated water transportation is usually accompanied by osmolyte accumulation to maintain a dynamic equilibrium of water potential (Wang *et al*. 2016). Our qRT-PCR results revealed that several *AQP*s, including *TONOPLAST INTRINSIC PROTEIN 1;2* (*OsTIP1;2*) and *PLASMA MEMBRANE INTRINSIC PROTEIN 1;3* (*OsPIP1;3*), were upregulated by ABA. *RePRP2OX* lines showed even higher transcript level of *OsPIP1;3* and *OsTIP1;2* compared to the WT under ABA treatment, whereas *RePRP*-RNAi lines showed a lower level of these two *AQP*s than the WT under ABA treatment (Fig. 6C–6D). These two AQPs were reported to facilitate water uptake or increase plant resistance against drought conditions (Li *et al*., 2008; Lian *et al*., 2004). Our western blot and proteomic analyses indicated that the protein levels of PIP;3 and TIP1;2, together with various PIPs and TIPs, showed more than 1.5-fold upregulation in the *RePRP2OX* line even in the absence of ABA (Fig. 6E; Table S7). We validated that OsPIP1;3 and OsTIP1 proteins were increased in level in *RePRP2OX* lines under control conditions, which resembled WT roots treated with ABA. In contrast, the ABA-mediated induction of TIP1;1/TIP1;2 seemed attenuated in *ReRPP*-RNAi lines (Fig. 6F). These results indicate that *RePRP*s are responsible for abiotic stress-mediated induction of AQPs that may facilitate water transportation in root cells during drought and elevate overall survivability. In summary, *RePRP2* is required for controlling root water equibrium by increasing AQP levels and osmolyte concentrations, which further enhances root resistance to water deprivation.

## Discussion

### *RePRP*s augment rice with drought resistance by enhancing water conservation mechanisms

Appropriate control of water loss and water uptake in response to drought is critical for rice production. In this work, we demonstrated that RePRPs play critical roles in establishing tolerance to osmotic stress and soil drying. We further explored the underlying mechanisms leading to the enhanced stress tolerance and found that RePRPs are crucial for mediating root architecture, water equilibrium and shoot stomatal conductance in response to drought.

Here we showed unequivocally that *RePRP*s, especially *RePRP2*, augmented drought resistance in plants by reducing water loss (Fig. 2, Fig 3, Fig 5, Fig S7–S10) and lowering root osmotic potential (Fig. 6 and Fig. S12). By promoting cell wall-bound POX synthesis and lignin/suberin biosynthesis, *RePRP*s facilitated the formation of water barriers and prevented water loss under extreme drought conditions. The increase in cell wall-bound POX activities found critical for lignin polymer assembly, and ABA can trigger suberin and lignin deposition in the wall of endodermis cells toward root tip regions (Lin and Kao, 2001a; Schreiber *et al*. 2005; Lee *et al*. 2013; Lin *et al*., 2016). As a result, the integrity of exodermis and endodermis (Casparian strips) will be increased, leading to decreased water loss under drought stress (Cruz *et al*. 1992). ABA treatment could not further increase lignin levels in our *RePR2OX* lines, likely due to lack of sufficient supply of lignin precursors because genes involved in lignin biosynthesis were only slightly upregulated in *RePRP2OX* (Fig. 5, Table S5). Also, ABA-induced lignin accumulation in roots was mainly accumulated in the vascular system, but *RePRP2OX* lines showed lignin accumulation in both the exodermis and vasculature (Fig. S9 and Fig. S10). The results suggest that ectopic expression of *RePRP* can specifically enhance root water conservation without disrupting overall lignin metabolism. Our study also suggested an *RePRP*-independent molecular mechanism that can trigger ABA-medaited lignin accumulation, H_2_O_2_ production and POX activation because the *RePRP*-RNAi lines acted similar to the WT when ABA was applied (Fig. 5E–5G).

Drought- or ABA-responsive genes can act redundantly to maintain proper stress resistance. For example, *PKABA1*, an ABA-induced protein kinase, is sufficient but not necessary for suppressing the expression of the TF *GAMyb* that triggers the expression of α-amylase (Zentella et al., 2002). Drought also triggers the expression of NACs and drought-responsive element binding proteins that enhance plant drought tolerance independent of ABA (Shinozaki and Yamaguchi-Shinozak, 2007). Future work on identifying unknown factors that act redundantly to RePRPs will be crutial for clarifying the roles of RePRPs in response to both ABA and drought.

*RePRP2OX* caused lower overall stomatal conductance (Fig 2), which is probably one of the main reasons for *RePRP2OX* plants consuming less water (Fig 3B and 3C). The disassembly of microtubules is required for ABA-mediated stomata closure (Dou *et al*., 2021). Our previous study indicated that *RePRP*s were essential for microtubule re-orientation (Hsiao *et al*., 2020). The decrease in stomata conductance observed in *RePRP2OX* was probably related to the function of *RePRP*s in proper microtubule formation. Also, in Arabidopsis, the development of stomata is regulated by cutin, which is similar to suberin in structure (Yang *et al*., 2022). Future studies of the potential involvement of *RePRP*-mediated lignin/suberin formation and microtubule orientation in rice stomatal development will shed light on improving rice water conservation.

With respect to water uptake, *RePRP2* enhanced drought tolerance by improving WUE in root. We also validated that RePRP2 is responsible for ABA-induced expression of the AQPs PIP1;3 and TIP1;2 (Fig. 6C–6F, Table S7). The specific increase in AQP levels in root may facilitate plant water movement under drought. The level of root PIP abundance and plant hydraulic conductance have been found highly related to leaf growth in a circadian oscillation-dependent manner (Caldeira *et al*. 2014; Maurel *et al*. 2015). *PIP1;2* and *PIP1;3* transcript levels were increased in upland rice that were generally more resistant to drought than lowland rice (Lian *et al*., 2006). Stress-inducible promoter-driven *PIP1;3* expression enhances rice drought resistance (Lian *et al*. 2004). Strawberry TIPs (FaTIPs) were upregulated in drought-tolerant cultivars under severe drought, which was associated with gradual stomata closure and prevention of water loss under dehydration (Merlaen *et al*., 2019). Overexpression of TIP1;2 could increase water transportation of *Xenopus* oocytes (Li *et al*., 2008). We also observed a decrease in root osmotic potential in *RePRP2OX* lines that was partly due to the accumulation of the osmolyte proline (Fig. 6A–6B, Fig. S12). Taken together, ectopic expression of *RePRP* is beneficial for improving rice water conservation in response to drought without many trade-offs on overall plant growth.

### Plausible mode of action of *RePRP*s under stress

*RePRP2OX* can upregulate several ABA-inducible *LEA* genes and many stress-responsive TFs (Fig. 4, Fig. S4 and S5, Table S3 and S4) that may play pivotal roles in stress responses. For example, overexpression of *DROUGHT RESPONSIVE ZINC FINGER PROTEIN 1* (*DRZ1*) enhanced rice seedling drought tolerance (Yuan et al., 2018). The group VII ERFs and their activators were found essential for regulating rice abiotic and biotic stress responses (Licausi *et al*., 2013). Our transcriptomic analyses also revealed that several group VII ERFs, including *ERF60*, *ERF61*, *ERF67* and *ERF68*, were induced by ABA in an *RePRP2*-dependent manner (Fig. S5). *HSF15/SPL7* was required to maintain reactive oxygen species (ROS) balance and rice stress responses (Hoang *et al*., 2019). *NAC95* is stress responsive and is involved in cell wall development (Shen et al., 2009; Sun *et al*., 2015). *NAC66* promoted rice stress tolerance by increasing the concentration of osmolytes such as sugar and proline (Yuan *et al*., 2019). *WRKY21* content was related to the expression of the NAD kinase *OsNADK1* that enhanced rice drought tolerance by maintaining cell redox balance (Wang et al., 2020). *WRKY76* was reported to increase the expression of abiotic stress-responsive genes such as *POX*s (Yokotani *et al*. 2013). Thus, *RePRP*-mediated physiological changes may be associated with these stress-responsive TFs. However, *RePRP2* does not contain any known TF domains in its amino acid sequence, so the promotion of *LEA* and TF expression must be mediated by unknown factor(s) that may physically interact with *RePRP2* or relies on the unique sequence features of RePRPs. All four RePRPs show highly repetitive PEPK motifs, with > 40% proline in amino acid composition (Tseng *et al*., 2013; Hsiao *et al.,* 2020). Proline disrupts the formation of α-helix and causes an intrinsically disordered protein structure (Sun *et al*. 2013). Proline-rich domains also serve as modulators of their environments (Reiersen and Rees, 2001), providing more efficient changes in protein structures as compared with most globular proteins. The pleiotropic regulatory functions of RePRPs may be related to intrinsically disordered properties.

### Multifaceted integrative roles of *RePRP*s and their potential involvements in multiple stress tolerance

As a group of intrinsically disordered proteins with the potential of interacting with multiple partners, RePRPs elicit multifaceted responses in rice leading to a high level of drought tolerance. In the shoot, *RePRPOX* causes reduced stomata conductance probably by reduced stomata aperture. In the root, water barriers such as lignin and suberin are formed in the vascular system and root peripherals to prevent water loss. However, at the same time, osmotic pressure in roots becomes more negative and movement of water in the plant tissues is facilitated by elevated levels of specific AQPs. The combination of these events reveals the versatile nature of these intrinsically disordered proteins. *RePRPOX* plants are able to consume much less water overall yet manage to have longer roots under moderate osmotic stress, which is likely related to the extraordinary tolerance to drought stress.

Although not investigated in this study, the genes upregulated by *RePRP2OX* but not ABA may, at least in part, be involved in biotic stress responses (Table S4). One report showed that expression of a transgene encoding a *Cajanus cajan* hybrid proline-rich protein may increase rice resistance to abiotic and biotic stress (Mellacheruvu *et al*. 2016), although the underlying mechanisms remain to be investigated. Our transcriptomic analyses indicated that genes upregulated by *RePRP2OX* are also involved in chitin catabolism and diterpenoid biosynthesis (Table S4), which suggests that *RePRP*s may also be involved in the defense response to biotic stresses. Future studies of the integrative roles of *RePRP*s will be valuable to improve rice resistance against threats of abiotic and biotic stresses. Furthermore, this group of unique stress-induced intrinsic disordered proteins may be an important target for technology development in enhancing stress resistance in cereals.

## Materials and methods

### Plant materials and growth conditions

Rice seeds with Japonica TNG67 background were sterilized with 2% sodium hypochlorite for 30 min, then washed thoroughly with distilled water. The washed seeds were germinated in water at 28 °C for 3 days. Uniformly germinated seeds were selected and cultivated in hydroponic culture by using 1/2 Kimura B solution (Hsu and Kao, 2003). The cultivated seedlings were grown at 28 °C at 90% relative humidity in 14-h light/10-h dark conditions with light intensity 200 µmol m^-2^ s^-1^.

### Abiotic stress treatments

For drought treatment in the hydroponic system, WT and transgenic rice plants (10 seedlings per line) were grown in the same tray for 2 weeks under 1/2 Kimura B solution. Two-week-old seedlings (three leafs) were exposed to 30% PEG in distilled water (PEG 6000; Merck) for 18-20 h. The treated seedlings were washed thoroughly to remove residual PEG on the roots and were recovered in fresh 1/2 Kimura B solution. Survival rates were evaluated on day 12 of recovery. For soil cultivation, 10-day-old seedlings grown in 1/2 Kimura solution were transferred to a 7.62-cm pot containing soil mixture (2:1 v/v peat moss and perlite, Jiffy) and saturated with water (190 g per pot). The seedlings were grown for an additional 21 days at 28°C under long-day conditions (14-h light/10-h dark, 200 μmol m^-2^ s^-1^). The 31-day-old seedlings were saturated with water and pots were placed on a dry tray for 4 days, then recovered by watering. Survival was calculated on days 9 and 15 of recovery.

### Generation of transgenic plants

The generation of *RePRP2.1*-overexpression lines and *RePRP*-RNAi and *RePRP2*-RNAi lines was previously described (Tseng *et al*., 2013). An abiotic stress-inducible *RePRP2.1* construct was obtained by fusing the *RePRP2.1* coding sequence to the *3XABRC321* promoter (Chen *et al*., 2015) in the pENTR vector (ThermoFisher, USA). The combined fragment was cloned into the pZP200 binary vector by LR recombination (ThermoFisher, USA). Transgenic rice lines were generated in the rice transformation laboratory at the Institute of Molecular Biology, Academia Sinica via *Agrobacterium*-mediated transformation as described (Hong *et al*., 2004).

### RNA sequencing and bioinformatic analysis

Ten-day-old rice seedlings grown in 1/2 Kimura solution under long day conditions (14-h light/10-h dark) were treated with or with 2 μM of ABA for 96 h, and root tips (1 cm from the top of roots) were collected for total RNA extraction. Total RNA were extracted by using TRIZOL reagent (ThermoFisher, USA) and the QIAGEN RNAeasy kit, following the manufacturers’ instructions. About 10 μg total RNA was used for mRNA purification followed by cDNA library preparation with the TruSeq Stranded mRNA kit (Illumina, USA). The prepared cDNA libraries were amplified by PCR and sequenced by using Illumina Hiseq 2500. About 70-100 million reads per sample were obtained and underwent transcriptomic analyses. Adapter removal was done on raw reads with Trimmomatic (Bolger *et al*., 2014); clipped reads shorter than 100 bp were dropped. According to the genome sequences and genome annotation from MSU Rice Genome Annotation Project Release 7 (Kawahara *et al*., 2013), passed reads were first mapped to the rice transcriptome and unmapped reads were then mapped to the rice genome. With all alignments transferred to the genome coordinate, normalized read counts of genes were computed by using the RACKJ toolkit (http://rackj.sourceforge.net/) and the trimmed mean of M-values (TMM) method (Robinson and Oshlack, 2010). The mapping statistics are shown in Table S9. Expressed genes defined as genes with > 1 reads per kilobase per million reads (RPKM) in at least one sample were considered expressed genes, and the criterion had to be fulfilled in all three biological replicates. Genes with fold change > 2 or < 0.5 in all three biological replicates as compared with the WT control were considered differentially expressed genes. For functional characterization of differentially expressed genes, Gene Ontology and Kyoto Encyclopedia of Genes and Genomes databases were used for function mapping by DAVID V 6.7 (Huang *et al*. 2009). Significantly enriched GO terms and KEGG pathways were defined as terms/pathways with p < 0.01 on Fisher’s exact test, using all expressed genes as a background reference.

### Proteomic analysis

Ten-day-old seedlings of the WT and *RePRP2OX* line were treated with or without ABA (2 μM) for 4 days. Root samples (1 cm from root tip) were collected and ground into fine powder with the aid of liquid nitrogen. The powder was extracted with a 10X volume of trichloroacetic acid (TCA) solution (10% TCA, 0.07% β-mercaptoethanol in acetone) and incubated for 45 min at -20°C. The precipitated proteins were centrifuged for 15 min at 35,000 g, and pellets were washed with a 10X volume of 0.07% β-mercaptoethanol in acetone and incubated for 1 h at -20°C. The washed samples were centrifuged for 15 min at 35,000 g. The pellets were then washed and centrifuged twice by adding a 10X volume of 0.07% β-mercaptoethanol in acetone without further incubation. The pellets were dried by SpeedVac for 30 min and dissolved in dissolution solution (8M urea, 50 mM Tris-HCl, pH 8.5) to a final concentration of 1-5 mg/mL. The samples were then labeled with isobaric tags for relative and absolute quantification (iTRAQ, 4-plex) kit, following the manufacturer’s instructions. The labeled samples were trypsin-digested and underwent untargeted mass spectrometry followed by a Mascot search (Matrix Science, USA) in the NCBI rice Protein RefSeq. The complete search results are shown in Table S10. The protein abundance ratio to the WT control was used to calculate the protein fold changes of AQPs.

### Quantitative RT-PCR analysis

Total RNA was isolated from 10-day old rice roots treated with or without ABA (2 μM) by using TRIzol reagent (ThermoFisher) according to the manufacturer’s instructions. First-strand cDNA was synthesized from 2 μg total RNA by using the SuperScript III first-strand synthesis system (ThermoFisher), following the manufacturer’s instructions. Specific primer pairs used for detection of are in Table S11. The transcript levels of each gene examined were normalized to those of internal controls (*Actin1* or *EF-1α*) within the samples.

### Root water usage measurement

Four plants (10-day-old seedlings) were transferred to a glass tube containing an equal volume of 1/2 Kimura B solution with or without 10% PEG. The tubes were sealed with parafilm to avoid water evaporating. To evaluate the water used by rice plants, the total tube weight was measured every day for 1 week from the beginning of experiments. The plant roots and shoots from day 0 and day 7 were collected separately to measure dry weight and assess the biomass increase. Then, the water use efficiency (WUE) of plant tissues was calculated as tissue biomass increase (mg)/water used (g).

### Root osmotic potential measurement

Two-week-old seedlings of WT, *RePRP2OX* and *RePRP*-RNAi lines were treated with 15% PEG 6000 solution for 24 h. The treated samples were carefully washed three times with de-ionized distilled water (ddH_2_O). Whole roots were collected and immediately frozen at -20℃. Root tissues were ground with a microfuge pestle in a microcentrifuge tube. The powder was thawed and centrifuged to remove insoluble cell debris and the cell sap was transferred to a clean new tube. The osmotic potential measurement was quantified using a Psyprosystem with C-52 sample chambers (Wescor) (Kumar *et al*., 2013).

### Lignin/suberin staining and microscopy observation

Root tissues (1 cm from root tip) of 10-day-old seedlings treated with or without 2 μM ABA were submerged in fixation solution (ethanol: acetic acid: formaldehyde = 90: 5: 5). The samples were then infiltrated by vacuum for 10 min twice before being embedded in 5% agar. Cross-sections (50-100 μm thickness) generated by Vibratome MicroSlicers DTK-1000 (Ted Pella, USA) and stored in fixation solution. For lignin staining, the cross-section samples were stained with acriflavine (Sigma-Aldrich, cat no. 01673) as described (Rocha *et al*., 2014). In brief, sections of plant tissues were stained with 1 mL freshly prepared 0.0025% acriflavine (in ddH_2_O) for 10 min with gentle shaking (50 rpm) under dark, 25°C. The stained sections were washed with 1 mL ddH_2_O three times and mounted in 75% glycerol before observation. Images were observed and captured under a Zeiss LSM 510 META confocal microscope with a 488 nm laser for excitation and 505 to 550 nm band pass filter for emission. For suberin staining, Fluorol Yellow 088 (Setareh Biotech, cat. No. 6689) was used for suberin as described (Brundrett *et al*., 1991; Landgraf *et al*., 2014). In brief, 0.01% Fluorol Yellow 088 was dissolved in PEG4000 by heating to 90°C for 1 h. The heated Fluorol Yellow 088 solution was then mixed with an equal volume of 90% glycerol before use. The sectioned tissues were submerged in 1 mL FY088/PEG400/glycerol solution in a 12-well plate. The samples were stained for 1 h at 50 rpm, 25°C and washed with ddH_2_O three times (1 mL each time). The washed samples were mounted in 75% glycerol and imaged with use of a Zeiss LSM510 META confocal microscope with excitation wavelength 405 nm for autofluorescence and 488 nm for Fluorol Yellow 088. The emission filter settings were 418 to 480 nm for autofluorescence and 524 to 594 nm for Fluorol Yellow 088.

### Hydrogen peroxide (H_2_O_2_) staining and peroxidase activity determination

3,3-diaminobenzidine (DAB) staining was used for H_2_O_2_ detection in roots (Daudi *et al*. 2012). DAB was dissolved in 10 mM Na_2_HPO_4_, pH 6.8 (1 mg/mL). Ten-day-old rice seedlings treated with or without 2 μM ABA for 4 days were submerged in DAB solution and infiltrated by vacuum three times (5 min each time). The infiltrated seedlings were incubated in the DAB solution for an additional 10 min. Stained root tips were stored in 70% ethanol and images were captured by dissection microscope (ZEISS Stemi 2000-C) coupled with a CCD camera. Cell wall-bound POX activity was detected using guaiacol as a substrate as described (Lin and Kao, 2001b). In brief, 0.1 g of root tips was ground into fine powder with the aid of liquid nitrogen. The frozen tissues were mixed with 5 mL potassium buffer (50 mM K_2_HPO_4_/KH_2_PO_4_, pH 5.8) and centrifuged for 10 min at 1000 xg, 4°C. After each centrifugation, 4 mL of potassium buffer was removed and re-supplemented with fresh potassium buffer to wash the tissues. After four centrifugations, 4 mL potassium buffer was removed and 4 mL of 1M NaCl was added to extract the cell wall-bound peroxidases. After 2 h of extraction at 30°C, the samples were centrifuged for 10 min at 1000 xg, 4°C and the supernatant was transferred to a clean new tube for POX activity detection. For each sample, 100 μL of extracted enzymes was mixed with 1 mL potassium buffer, 1 mL of 21.6 mM guaiacol and 39 mM H_2_O_2_ and mixed by vortexing. A 200-μL amount of the mixture was immediately added to a quartz ELISA reader and the absorbance at 470 nm was detected. One unit of POX activity was defined as 1 mmol of tetraguaiacol produced per minute.

### Lignin extraction and lignin content determination

The acetyl bromide method was used for lignin extraction and determination (Hatfield *et al*. 1999; Kapp *et al*. 2015). In brief, 80 to 100 mg of root tip tissues was ground into fine powder in liquid nitrogen. The powder was washed three times with 95% EtOH, then three times with EtOH: hexane (1:2) mixture. To remove starch, the washed samples were incubated with 90% DMSO overnight with gentle shaking. The de-starched samples were washed six times with 70% EtOH, followed by a 1-mL acetone wash. The washed powders were air-dried and defined as alcohol insoluble residues (AIRs). About 5 mg of AIR was added to screw-cap glass vials and mixed with 1 mL acetyl bromide solution (in 25% glacial acetic acid) by vortexing. The mixtures were incubated for 1 h at 70°C in a water bath with gentle swirling. After incubation, the samples were cooled to room temperature before the addition of 0.9 mL of 2N NaOH, 0.1 mL of 7.5 M hydroxylamine hydrochloride and 3 mL glacial acetic acid. The stabilized samples were diluted 10 times by the addition of glacial acetic acid before detection by Gen5 ELISA reader using Black quartz microplate (Hellma), with detection wave length set at 280 nm. The extinction coefficient for calculation was set to 17.57, as in a previous study (Foster *et al*. 2010).

### Stomatal conductance measurement and stomata morphology observation with scanning electron microscopy (SEM)

Two-week-old rice plants were grown in pots and cultivated for an additional 2 months in the greenhouse. The pots were placed in a growth chamber for 1 week under long day conditions (14-h light/10-h dark, light intensity: 300 μmol/m^2^s) before measuring stomatal conductance. The CO_2_ concentration in air was 400 to 450 μmol/mol. Li-COR 6800 (Li-COR, USA) was used to measure stomatal conductance. The configuration of Li-COR 6800 was set as follows unless otherwise noted: flow: 500 μmol/s, area: 3 cm^2^, lamp: 1200 μmol/m^2^s, with red: blue light set to 9:1. For observation of leaf stomatal density and morphology, 2-week-old rice leaves were dissected and loaded onto stubs. The samples were frozen by liquid nitrogen slush and transferred to a sample preparation chamber at -160℃. After 5 min of incubation, the temperature was raised to -85℃ and the samples were sublimed for 15 min. After coating with Pt at -130℃, the samples were then transferred to cryo stage in the SEM chamber and stomatal morphology was observed at -160℃ by using a cryo scanning electron microscope (Quanta 200 SEM/Quorum Cryo System PP2000TR, FEI, USA) with 20 KV.

### Root protein extraction and western blotting

Ten-day-old rice seedlings grown in 1/2 Kimura solution were transferred in fresh 1/2 Kimura solution with or without 2 μM of ABA and grown for an additional 96 h. Whole root tissues were collected ground into fine powder with the aid of liquid nitrogen. About 100 mg tissue powder was extracted with 1 mL protein extraction buffer (50 mM Tris pH 8.0, 10 mM NaCl, 1% SDS, 0.1 mM dithiothreitol, 20 μM PMSF, 0.5% β-mercaptoethanol, 1X proteinase inhibitor from Roche) and centrifuged for 10 min at 4°C. The supernatant was transferred to a clean new tube and protein concentration was quantified with the DC Protein Assay kit (Bio-Rad). SDS-PAGE and western blot analysis were conducted as described (Tseng et al., 2013). The primary antibody against rice PIP1;3 (Agrisera, AS09 504) and *Arabidopsis thaliana/Raphanus sativus* TIP1;1/TIP1;2 (Agrisera, AS09 493) were used for AQP protein detection, with a dilution ratio of 1:5000. The secondary antibody (Anti-Rabbit IgG [Goat], HRP-Labeled, PerkinElmer NEF812001EA) was 1:10000 in 5% TBST-milk.

### Determination of root proline content

Proline isolation and measurement protocols were based on previous studies (Bates et al. 1973; Verslues et al., 2010), with minor modifications. Ten-day-old rice seedlings were treated with or without ABA (2 μM) for 24 h. Whole roots were collected and ground into fine powder with the aid of liquid nitrogen. About 100 mg of sample was extracted with 840 μL ddH_2_O and centrifuged to remove cell debris. For each sample, 200 μL supernatant was added to a clean tube and 300 μL ddH_2_O was added. The sample was then sequentially mixed with 500 μL acetic acid and 500 μL Chinard’s ninhydrin solution. The mixed sample was put into boiling water and reacted for 1 h. After reaction, the sample was cooled to room temperature and mixed with 1500 μL toluene by vortexing. The mixed sample was incubated for 15 min to separate the toluene and water layers. After incubation, 200 μL toluene layer was added to a quartz ELIZA reader plate and OD_595_ nm was measured. A serial dilution of proline was used as standard curves of each experiment.

### Root water uptake assay

Fresh weight of 10-day-old rice seedlings grown in 1/2 Kimura solution was measured and defined as 100% water content. The measured plants were dried on tissue paper for 3 h. After drying, the leaves were spread with Vaseline to prevent transpiration for aerial parts. The fresh weight before and after Vaseline spreading was recorded. The Vaseline-sealed plants were then incubated in glass beakers containing 100 mL distilled water and their fresh weight was recorded until 6 h after rehydration. The rehydrated plants were collected and dried at 50°C for 2 days and the plant dry weight was recorded and defined as 0% of water content.

### β-Glucuronidase activity staining

For GUS activity staining, we used transgenic rice harboring the *ProRePRP2.1::GUS* cassette, as described (Tseng *et al*., 2013). Ten-day-old rice seedlings grown in 1/2 Kimura solution were treated with or without 2 μM ABA in fresh 1/2 Kimura solution and grown for an additional 24 h before GUS activity staining as described (Jefferson et al., 1987), with slight modification. In brief, seedlings were submerged in 90% ice-cold acetone and incubated for 20 min at room temperature. After incubation, seedlings were transferred to a clean new tube containing GUS staining buffer (0.2% Triton X-100, 50 mM NaH_2_PO_4_, 2 mM potassium ferrocyanide, 2 mM potassium ferricyanide, pH 7.2) and incubated for 5 min on ice. The staining buffer was then removed and changed to fresh staining buffer containing 2 mM X-GlucA (Cyrus bioscience, Taiwan). The samples were vacuum-infiltrated for 5 min and transferred to a 37℃ dark chamber. After 4 h of incubation, the samples were transferred to clean tubes containing 70% EtOH and incubated for 16 h at 50℃. The stained leaf tissues were sliced and the GUS signal localization were observed by using a Zeiss Axio Observer Z1 motorized inverted fluorescence microscope under bright field.

### Elemental analyses

The elemental analysis procedures were as described (Lin *et al*., 2009). Root tissues were harvested separately and washed with 10 mm CaCl_2_ for 20 min and H_2_O for 10 min. Samples were dried at 70°C for 3 days. Dried root samples (about 20 mg) from 25 plants were transferred into a Teflon vessel and digested with 2 mL of 65% HNO_3_ and 0.5 mL H_2_O_2_ (Suprapur; Merck, Darmstadt, Germany) in a MarsXpress microwave digestion system (CEM, Matthews, NC, USA). Tomato leaves (SRM-1573a) from the National Institute of Standards and Technology (Gaithersburg, MD, USA) were used as a reference. The volume of solution after digestion was adjusted to 8 mL with H_2_O and filtered through a 0.45-µm membrane filter. The concentrations of K, Na, P, Fe, and Ca in digested samples were analyzed by inductively coupled plasma-optical emission spectrometry (ICP-OES) (OPTIMA 5300; Perkin-Elmer, Wellesley, MA, USA). The concentration of each element was determined in three biological replicates.

#### Accession numbers

*RePRP2.1* (LOC_Os07g23660), *PIP1;3* (LOC_Os02g57720), *TIP1;2* (LOC_Os01g74450), *OsActin1* (LOC_Os03g50885), *EF-1α* (LOC_Os03g08020), *OsHSF15* (LOC_Os05g45410), *ONAC066* (LOC_Os03g56580), *POX51*-like (LOC_Os03g55410), *Prx86* (LOC_Os06g35520), *Prx90* (LOC_Os06g48030).

## Acknowledgments

We thank Dr. Wann-Neng Jane in the Cell Biology Core, Institute of Plant and Microbial Biology, Academia Sinica, for kind help in establishing the Cryo-SEM system; Dr. Wen-Dar Lin in the Bioinformatics Core Lab, Institute of Plant and Microbial Biology, Academia Sinica, for help with transcriptomic analyses; and Dr. Chuan-Chih Hsu in the Proteomics Core Lab, Institute of Plant and Microbial Biology, Academia Sinica, for help with identifying RePRP2-interacting proteins. We also thank them for helpful discussions in conducting related experiments.

## Supplemental data

**Figure S1. Drought tolerance of individual *RePRP2OX* and *RePRP*-RNAi lines.**

**Figure S2. Transcript levels of *RePRP2* under ABA treatment.**

**Figure S3. *RePRP2* is necessary for ABA-mediated induction of transcription factors.**

**Figure S4. Mapping results of reads within *HSF15* and *ONAC066* loci.**

**Figure S5. *RePRP2* is necessary for ABA-mediated induction of *LEA* genes.**

**Figure S6. Close-up view of suberin fluorescence under control conditions.**

**Figure S7. Close-up view of suberin fluorescence under ABA treatment.**

**Figure S8. Close-up view of lignin fluorescence under control conditions.**

**Figure S9. Close-up view of lignin fluorescence under ABA treatment.**

**Figure S10. *RePRP2OX* leads to increased root WUE and decreased overall water use.**

**Figure S11. Element concentrations was comparable among WT and *RePRP2OX* and *RePRP*-RNAi lines.**

**Figure S12. *RePRP2* is essential for ABA-induced expression of genes involved in proline metabolism.**

**Table S1. Genes upregulated by ABA or *RePRP2* overexpression.**

**Table S2. Actual RPKM values of transcription factors showed ABA- and *RePRP2*-dependent differential expression.**

**Table S3. Actual RPKM values of *LEA* gene transcripts; asterisks indicate the LEAs showing ABA- and *RePRP2*-dependence.**

**Table S4. GO and KEGG analyses of genes upregulated by ABA and/or *RePRP2* overexpression.**

**Table S5. RPKM values of lignin biosynthesis genes from the root tip transcriptome.**

**Table S6. Actual RPKM values of cell-wall-bound peroxidases showing ABA- and *RePRP2*- dependence.**

**Table S7. Relative protein levels of identified rice aquaporins under ABA treatment or *RePRP2* overexpression.**

**Table S8. Actual RPKM values of proline biosynthesis gene transcripts.**

**Table S9. Mapping statistics of RNA sequencing.**

**Table S10. Proteomic analysis of WT and *RePRP2OX* line. Table S11. Primers used for qRT-PCR.**

**Large dataset:** RNA sequencing files were deposited in NCBI BioProject (Project ID: PRJNA763046)

## Supplemental figure legends

**Figure S1. Drought tolerance of individual *RePRP2OX* and *RePRP2*-RNAi lines.**

**A** The three independent *RePRP2OX* lines showed increased survival after drought (30% PEG), compared to the wild type (WT). Three-week-old rice plants grown in ½ Kimura solution were treated with 30% PEG for 20 h, followed by recovery in ½ Kimura solution. The survival rate was calculated on day 15 after recovery. Data were analyzed by one-way ANOVA with Tukey’s multiple comparison test. Data are means ± SD from four biological repeats, with 30 seedlings per line in each repeat. Different superscript letters indicate significant mean difference (p < 0.05).

**B** *RePRP2OX* lines showed increased survival rate after 25% PEG treatment. Three-week-old seedlings were treated with 25% PEG for 3 days; the survival rates of each experiment were calculated after 15 days of recovery from PEG treatment.

**C** *RePRP2OX* lines showed increased survival rate after recovering from soil drought conditions. One-month-old soil-grown rice plants were treated with 4 days of dehydration before re-watering. The survival rate was calculated at day 15 after recovery. The horizontal line, boxes and whiskers are mean ± SD from 40 individuals At least four independent experiments were conducted and all showed similar results. Data were analyzed by one-way ANOVA with Tukey’s multiple comparison test. Different superscript letters indicate significant mean difference (p < 0.05).

**Figure S2. Transcript levels of RePRP2 under ABA treatment.**

**A** Independent *RePRP2OX* lines showed elevated *RePRP2* transcript levels without ABA treatment, but independent *RePRP*-RNAi lines showed suppressed ABA-induced *RePRP2* expression. Ten-day-old seedlings grown in ½ Kimura solution were treated with 2 μM ABA (in ½ Kimura solution) for the time indicated on the x-axis; whole root samples were collected for total RNA extraction and qRT-PCR. qRT-PCR using primers that amplified the *RePRP2* coding sequence was used to examine the transcript levels in independent *RePRP2OX* and *RePRP*-RNAi lines.

**B C***RePRP2.1* was induced by ABA in both root and shoot, but the transcript level was lower in shoot than root. qRT-PCR was used to validate the mRNA level of *RePRP2.1*. Ten-day-old seedlings grown in ½ Kimura solution were treated with or without 20 μM ABA for an additional 96 h. Data are mean ± SD. Data were analyzed by one-way ANOVA with Tukey’s multiple comparison test. Different superscript letters indicate significant mean difference (p < 0.05).

**Figure S3. *RePRP2OX* leads to increased root WUE and decreased overall water use.**

**A** Root WUE in TNG67 WT, *RePRP2OX* and *RePRP*-RNAi lines. Data were analyzed by one-way ANOVA with Tukey’s multiple comparison test. Different superscript letters indicate significant mean difference (p < 0.05).

**B** Shoot WUE in TNG67 WT, *RePRP2OX* and *RePRP*-RNAi lines. Data were analyzed by one-way ANOVA with Tukey’s multiple comparison test. Different superscript letters indicate significant mean difference (p < 0.05).

**C** Overall WUE in *RePRP2OX* and *RePRP*-RNAi lines. Data were analyzed by one-way ANOVA with Tukey’s multiple comparison test. Different superscript letters indicate significant mean difference (p < 0.05).

**D** *RePRP2OX* lines used less water, and *RePRP*-RNAi lines consumed similar amount of water as the WT. Data were analyzed by one-way ANOVA with Tukey’s multiple comparison test. Different superscript letters indicate significant mean difference (p < 0.05).

Five 10-d-old seedlings grown in ½ Kimura solution (14-h light/10-h dark) were transferred to glass tubes containing 85 mL ddH_2_O and sealed with Parafilm. Plants were grown for an additional 7 days under the same growth chamber, and water volume consumed was measured at day 7. Root and shoot tissues were collected separately and dried in a 50°C oven for 14 days to collect dry weight data from day 0 and day 7.

**Figure S4. *RePRP2* is necessary for ABA-mediated induction of transcription factors.**

The ABA-upregulated transcription factor genes (≥2-fold increase vs the WT control) were mostly upregulated in *RePRP2OX* lines under control conditions. Data mean fold change in expression from three biological repeats of RNA sequencing. Asterisks indicate the transcription factors reported to be involved in regulating rice drought resistance. The “red-red-blue” pattern indicates the expression of these genes were ABA and *RePRP2*-dependent.

**Figure S5. Mapping results of reads within *HSF15* and *ONAC066* loci.**

Integrative Genome Viewer (IGV) results were used to visualize the read mapping of each independent repeats. The read count scale for each gene (0 to 250 for *HSF15* and 0 to 100 for *ONAC066*) were set to be equal for easier comparison between samples. Both *HSF15* and *ONAC066* showed increased read count in the WT treated with ABA and *RePRP2OX* under control conditions. The mapping results of *RePRP*-RNAi line indicate that ABA application failed to induce the expression of *HSF15* and *ONAC066* when *RePRP* levels were decreased.

**Figure S6. *RePRP2* is necessary for ABA-mediated induction of *LEA* genes.**

The ABA-upregulated *LEA* genes (≥2-fold increase in expression vs the WT) were mostly upregulated in *RePRP2OX* lines under control conditions. Data are mean fold change in expression from three biological repeats of root transcriptome. Asterisks indicate the ABA- and *RePRP2*-dependent LEAs. DHN: dehydrin; SMP: seed maturation protein. The “red-red-blue” pattern indicates the expression of these genes were ABA- and *RePRP2*-dependent.

**Figure S7. Close-up of suberin fluorescence under control conditions.**

Suberin fluorescence appeared at both exodermis and xylem but was enriched in xylem area. One representative of at least 10 images. Ten-day-old rice seedlings were grown in ½ Kimura solution and transferred to fresh ½ Kimura solution for additional 96 h. Root sections were cut at 1 cm from root tips and Fluorol Yellow 088 staining was used to visualize suberin accumulation.

**Figure S8. Close-up of suberin fluorescence under ABA treatment.**

Suberin fluorescence was enriched in xylem area after ABA treatment versus the control condition. One representative of at least 10 images. Ten-day-old rice seedlings were grown in ½ Kimura solution and transferred to fresh ½ Kimura solution containing 2 μM ABA for an additional 96 h. Root sections were cut at 1 cm from root tips and Fluorol Yellow 088 staining was used to visualize suberin accumulation.

**Figure S9. Close-up of lignin fluorescence under control condition.**

Lignin fluorescence appeared at both exodermis and xylem but was enriched in xylem area. One representative of at least 10 images. Ten-day-old rice seedlings were grown in ½ Kimura solution and transferred to fresh ½ Kimura solution for additional 96 hours. Root sections were cut at 1cm from root tips and Acriflavine staining was used to visualize lignin accumulation.

**Figure S10. Close-up view of lignin fluorescence under ABA treatment.**

Lignin fluorescence appeared at both exodermis and xylem but was enriched in xylem area. One representative of at least 10 images. Ten-day-old rice seedlings were grown in ½ Kimura solution and transferred to fresh ½ Kimura solution containing 2 μM ABA for an additional 96 h. Root sections were cut at 1 cm from root tips and Acriflavine staining was used to visualize lignin accumulation.

**Figure S11. Element concentrations were comparable among WT and *RePRP2OX* and *RePRP*-RNAi lines.**

**A** Potassium content was comparable between WT, *RePRP2OX* and *RePRP*-RNAi lines.

**B** Sodium content was comparable between WT, *RePRP2OX* and *RePRP*-RNAi lines.

**C** Phosphorus content was comparable between WT, *RePRP2OX* and *RePRP*-RNAi lines..

**D** Iron content was comparable between WT, *RePRP2OX* and *RePRP*-RNAi lines.

**E** Calcium concentration was similar between WT, *RePRP2OX* and RePRP-RNAi root extracts. Ten-day-old rice seedlings grown in ½ Kimura solution were treated with or without 2 μM of ABA in ½ Kimura solution for 24 h. Whole root tissues were collected for ICP-OES analysis.

Data are mean ± SD from three technical repeats. Three biological repeats were conducted and all showed similar results.

**A** qRT-PCR of *P5CS* level in WT, *RePRP2OX* and *RePRP*-RNAi lines treated with ABA (2 μM). *RePRP*-RNAi line showed decreased *P5CS* level under control condition and the effect of ABA was attenuated.

**B** *RePRP2OX* line showed increased level of *ProDH* under control condition. The transcript level of *ProDH* was repressed by ABA application.

**C** The transcript levels of *P5CR*, the gene responsible for proline catabolism, were also highly dependent on the level of *RePRP2* and the presence of ABA. The level of P5CR was increased in *RePRP2OX* under control condition, similar to WT treated with ABA. *RePRP*-RNAi line showed decreased *P5CS* level under control condition and the effect of ABA was attenuated. Ten-day-old seedlings were treated with or without ABA (2 μM) for 96 h before root tissue collection (1 cm from root tip) and total RNA extraction.

Data are mean ± SD from three technical repeats. At least two biological repeats were conducted and all showed similar results. Data were analyzed by one-way ANOVA with Tukey’s multiple comparison test. Different superscript letters indicate significant mean difference (p < 0.05).

